# NEK7 accelerates NLRP3 inflammasome activation

**DOI:** 10.1101/2025.09.26.678733

**Authors:** Svenja Wöhrle, Tamara Ćiković, Clara Dufossez, Emilia Neuwirt, Elena Puma, Franziska Kraatz, Anna Kostina, Oliver Gorka, Clemens Kreutz, Christina J. Groß, Olaf Groß

**Affiliations:** Institute of Neuropathology, Faculty of Medicine, Medical Center, University of Freiburg, Freiburg, Germany; Faculty of Biology, University of Freiburg, Freiburg, Germany; Signalling Research Centres BIOSS and CIBSS, University of Freiburg, Freiburg, Germany; Institute of Medical Biometry and Statistics, Faculty of Medicine, Medical Center, University of Freiburg, Freiburg, Germany

## Abstract

The NLRP3 inflammasome is a major driver of immunopathology, making it a sought-after drug target. In spite of two decades of intense research, its precise activation mechanism remains elusive, impeding inhibitor design. NEK7 was reported as essential for NLRP3 activation, and several newly identified inhibitors were suggested to act by interfering with their interaction. Here we report that NEK7 accelerates, but is in principle dispensable for NLRP3 activation. The onset of inflammasome activation was unaltered in the absence of NEK7, yet the rate of cells to undergo inflammasome formation and subsequent pyroptosis was approximately 4-fold reduced. Therefore, therapeutic targeting of the NEK7-NLRP3 interaction might have an incomplete effect, which should be considered for drug development. We confirmed entrectinib as a NEK7-dependent inhibitor, while other published compounds turned out not to rely on it. Our results support two possible scenarios for the role of NEK7 in NLRP3 activation: either, NEK7 accelerates one unique pathway of NLRP3 activation, or it is essential for a fast pathway, while being dispensable for a second, slower mode of NLRP3 activation.

## Introduction

Inflammasomes are danger sensing platforms of innate immunity found in the cytoplasm of immune and epithelial cells. Through their constituent protease caspase-1, they control the maturation and unconventional secretion of leaderless interleukin-1 family cytokines, and lytic cell death through pyroptosis^1,2^. Of the different intracellular inflammasome-nucleating receptors, NLRP3 stands out by its propensity to drive immunopathology in numerous common acquired, as well as rare hereditary diseases. Its activation mechanism is unclear, and the development of specific tolerable inhibitors has been challenging^3–5^. Its numerous structurally divers activators of microbial or sterile endogenous or environmental origin have common effects on the cell that are thought to simultaneously trigger NLRP3 indirectly ^6–8^. These effects include plasma membrane, organellar, and metabolic stress, and cause potassium (K^+^) efflux ^9–13^, the generation of specific phospholipids on recycling endosome – trans-Golgi network (TGN) vesicles^14,15^, and inhibition of basic mitochondrial functions such as ATP production^16,28^. Although inhibiting K^+^ efflux, preventing NLRP3 recruitment to vesicle phospholipids, scavenging reactive oxygen species (ROS), or restoring oxidative phosphorylation (OXPHOS) blocks NLRP3 activation, how exactly these events are sensed by NLRP3 and cause activation is unknown. Posttranslational modifications also affect NLRP3, with alterations thereof generally perceived as prerequisite but in itself insufficient for activation^17–19^. Priming through cell surface receptors engaging NF-κB (typically through TLR4 activation with lipopolysaccharide [LPS]) is required for proIL-1β production, but also affects NLRP3 expression and posttranslational modification status (the latter referred to as *posttranslational priming* or *licensing*). Structural and functional insights suggest that the actual activation is a multi-step process involving its intrinsic ATPase activity, and the formation of both inactive and active NLRP3 multimers^20–22^. NEK7 belongs to a family of 11 NIMA-related kinases that are involved in microtubule organization and thereby, mitosis, meiosis, or ciliary biology^23^. NEK7 was reported to also be essential for NLRP3 activation, although the specific signal it relates to NLRP3 remained unclear^24,25,52^. However, emerging evidence hinted at a less pronounced requirement for NEK7^26,27^. We therefore reevaluated its impact on NLRP3 activation.

## Results

### NLRP3 inflammasome activation in the absence of NEK7

In previous work, we observed residual signals of caspase-1 and IL-1β maturation and release in NEK7-deficient murine macrophages, while in cells from *Nlrp3^-/-^* mice, these are typically undetectable^12,28^. We therefore directly compared NEK7^52^ and NLRP3-deficient^29^ murine bone marrow-derived macrophages (BMDMs) differentiated from bone marrow progenitors (Figure 1A-D). A clear and substantial residual signal for released mature IL-1β was observed in *Nek7^-/-^* cells primed with LPS and treated with four different commonly used NLRP3 activators (the K^+^/H^+^ ionophore nigericin^13^, extracellular ATP triggering the receptor P2X_7_^13^, the small molecule CL097^28^, and gout-related monosodium urate [MSU] crystals^29^, while no signal was detectable from *Nlrp3^-/-^* cells by ELISA (Figure 1A). Determining caspase-1 cleavage and release from the same samples by immunoblotting confirmed clearly-detectable inflammasome activation in the absence of NEK7 (Figure 1B). The degree of remaining IL-1β production in the absence of NEK7 was in the range of 20-40% of the signal detected from wildtype cells (Figure 1C). In contrast and as expected, cells from both *Nek7^-/-^* and *Nlrp3^-/-^* mice responded like wildtype cells to transfected poly(dA:dT) DNA as an AIM2 inflammasome activator (Figure 1A-C). Another hall mark of inflammasome activation are large, insoluble aggregates referred to as specks formed by the inflammasome adapter protein ASC that connects the different cytoplasmic receptor proteins to caspase-1. ASC specks were also still observed in NEK7 knockout cells, but were absent in NLRP3-deficient cells upon nigericin treatment (Figure 1D). Similar results were obtained using human monocytic THP-1 cells, with cells deficient in NEK7 displaying substantial residual inflammasome activity not observed in the absence of NLRP3 (Figure 1E-G). Therefore, although NEK7 clearly has some influence over NLRP3, inflammasome activation is essentially still intact in its absence.

**Figure 1:**
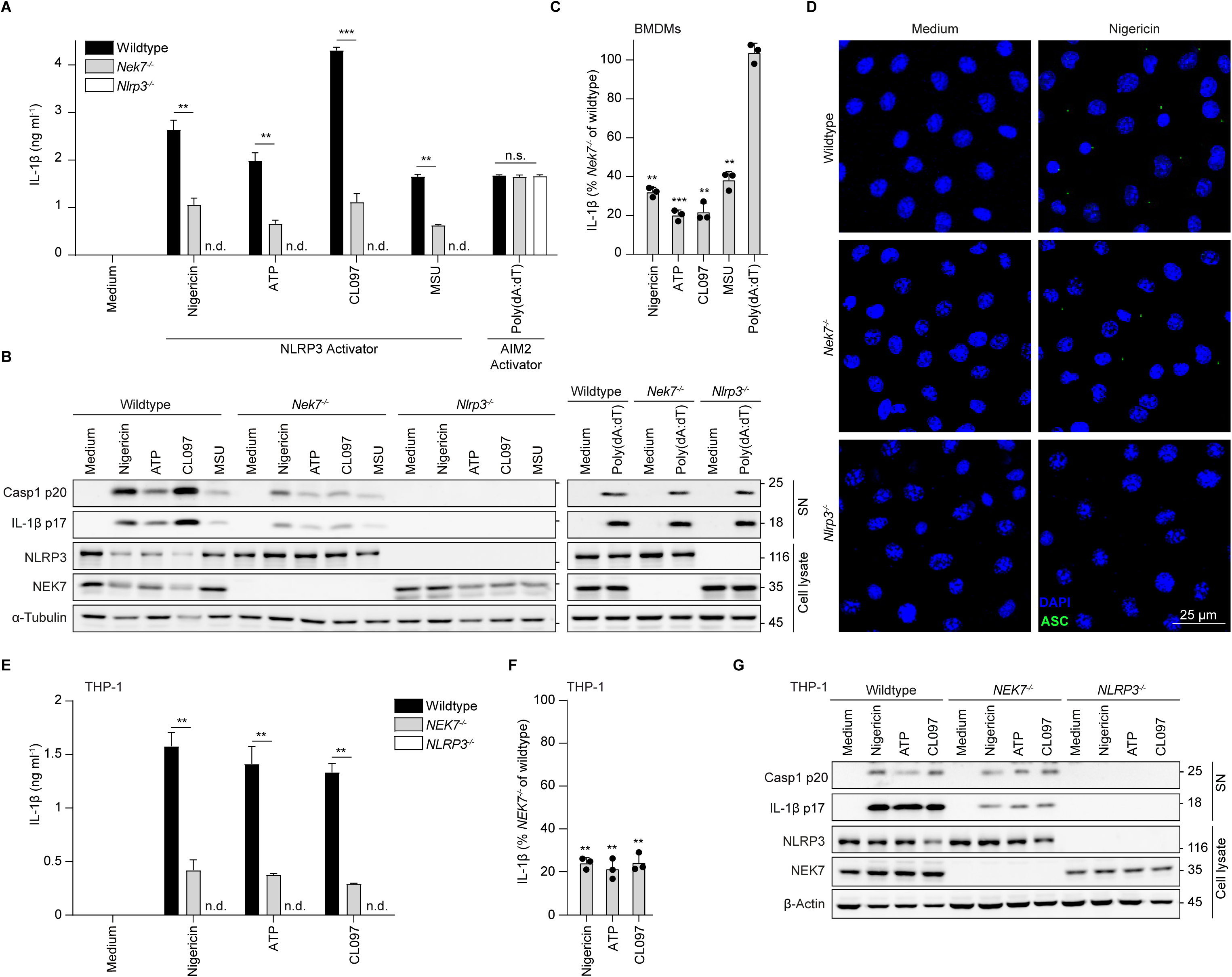
Residual inflammasome activation in Nek7 but not Nlrp3 knockout cells. (**A-C**) BMDMs from wildtype, *Nek7^-/-^* and *Nlrp3^-/-^* mice were primed with 20 ng ml^-1^ LPS and subsequently stimulated with 5 µM nigericin, 5 mM ATP, or 100 µM CL097 for 1.5 h or with 300 µg ml^-1^ MSU or 1 µg ml^-1^ poly(dA:dT) for 5 h. Release of IL-1β was determined from cell-free supernatants by ELISA, N = 3 technical replicates (A). (B) Immunoblot analysis of cell lysates and cell-free supernatants of LPS-primed wildtype, *Nek7^-/-^* and *Nlrp3^-/-^* BMDMs. (C) Residual inflammasome activation in *Nek7^-/-^* cells is depicted as percentage relative to wildtype. Each data point is derived by calculating the ratio of the mean of three technical replicates per genotype. Data of three independent experiments, one depicted in (A). Every dot is representative for an independent experiment, N = 3 biological replicates. (**D**) Confocal microscopy of LPS-primed BMDMs from wildtype, *Nek7^-/-^* and *Nlrp3^-/-^* mice were stimulated with 5 µM nigericin for 0.75 h. Hoechst 33342 (blue) localizes with the nuclei, ASC specks were stained with an anti-ASC antibody (green). Scale bar represents 25 µm. Images are representative for 2-3 images analyzed per condition depicting about 10-15 cells representative for 50,000 cells per well. (**E-G**) Wildtype, *NEK7^-/-^* and *NLRP3^-/-^* THP-1 cells were treated with PMA (200 ng ml^−1^) for 3 h, washed, and left at 37 °C, 5 % CO_2_ overnight. Subsequent the cells were primed with 150 ng ml^-1^ LPS and stimulated with 5 µM nigericin, 5 mM ATP or 100 µM CL097 for 2 h. IL-1β (N = 3 technical replicates) (E). Residual inflammsome activation (N = 3 biological replicates) (F) and immunoblotting (G) were measured and analyzed as in (A-C). ELISA, LDH, Immunoblot and microscopy data are representative of at least three independent experiments. For statistical analysis, METs were performed. Data are depicted as mean ±SD. Significance levels are denoted as *p < 0.05, **p < 0.01, ***p < 0.001, and ns for not significant.

### Priming does not affect the influence of NEK7 on NLRP3

In order to examine a potential influence of priming on NEK7-dependency, we began by comparing different pattern recognition receptor ligands (LPS for TLR4, Pam3CSK4 for TLR2, R848 for TLR7, and Zymosan primarily for Dectin-1, Figure 2A-D). Expectably, the different priming agents somewhat varied in the degree of priming as monitored by the induced amounts of proIL-1β and NLRP3, as well as of the Golgi-dependent cytokines TNF and IL-6 (Figure 2A, B). Although this reflected in also somewhat variable amounts of IL-1β released upon subsequent NLRP3 activation, the identity of the priming agent did not affect the degree of NEK7-dependency (Figure 2C, D). We next tested whether the intensity of priming has an effect, titrating two different TLR ligands, LPS and Pam3CSK4 (Figure 2E-I and Figure S1A, B). As expected, this again affected the absolute amounts of proIL-1β, TNF, and IL-6, as well as NLRP3 expression (Figure 2E and S1A, B). However, priming intensity also had no influence on the degree of dependency of mature IL-1β release on the presence of NEK7 (Figure 2F-I). Furthermore, priming intensity and choice of stimulus also did it not affect the degree of NEK7-dependency for pyroptosis, as quantified by determining lactate dehydrogenase (LDH) enzymatic activity in the supernatant from the same samples (Figure 2C, D, and F-I). Similar results were obtained for the duration of priming before inflammasome activation, which also influenced the total amounts of pro and mature IL-1β produced, but not the degree of NEK7-dependency (Figure 2J, K). This suggests that differences in the specific priming protocol cannot account for the variable degrees of NEK7-dependency observed.

**Figure 2:**
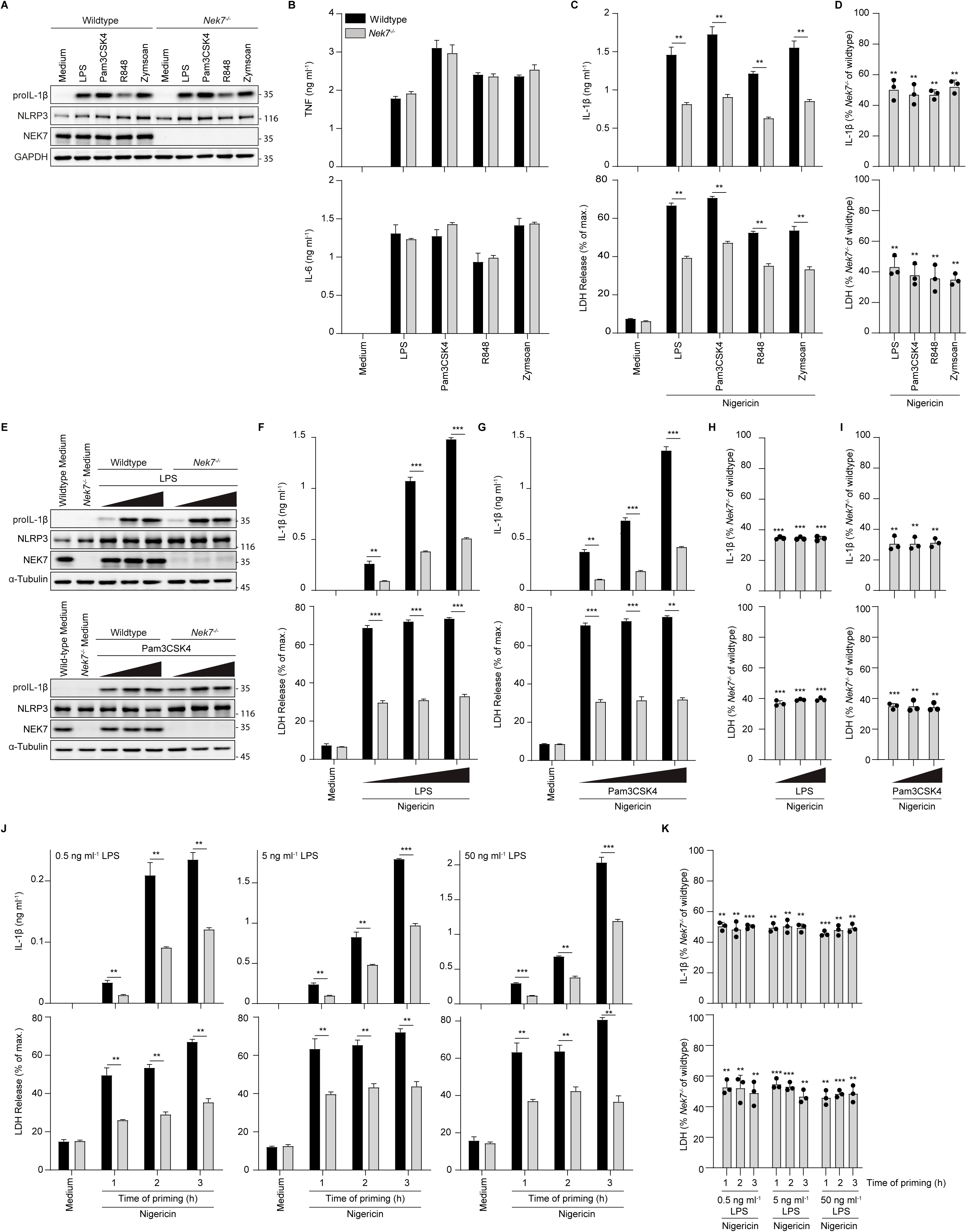
Priming does not affect the influence of NEK7 on NLRP3. (**A-D**) BMDMs from wildtype and *Nek7^-/-^* mice were primed with 1 µg ml^-1^ Pam3CSK4, 50 ng ml^-1^ LPS, 27 µM R848 or 1.5 µg ml^-1^ Zymosan for 3 h, harvested (A and B) or stimulated with 5 µM nigericin for 2 h (C, D). Production of proIL-1β (A), TNF and IL-6 (B) after priming was analysed by immunoblotting (A) or ELISA (B). Release of IL-1β and LDH after nigericin stimulation was determined from cell-free supernatants by ELISA and a colorimetric assay, respectively (C). N = 3 technical replicates. (D) Residual inflammasome activation in *Nek7^-/-^* cells is depicted as percentage relative to wildtype. Each data point is derived by calculating the ratio of the mean of three technical replicates per genotype. Data of three independent experiments, one depicted in (C). Every dot is representative for an independent experiment, N = 3 biological replicates. (**E-I**) BMDMs from wildtype and *Nek7^-/-^* mice were primed with increasing concentration of LPS (0.4, 2, 10, 50 ng ml^-1^) or Pam3CSK4 (0.125, 0.25, 0.5, 1 µg ml^-1^) for 3 h and harvested for immunoblotting (E) or subsequently stimulated with 5 µM nigericin for 2 h (F-I). IL-1β and LDH secretion and proIL-1β production were determined as in A-C and ratios calculated as in (D). (**J-K**) BMDMs from wildtype and *Nek7^-/-^*mice were primed with increasing concentrations of LPS (0.5 - 50 ng ml^-1^) for 1-3 h and stimulated with 5 µM nigericin for 2 h. Release of IL-1β and LDH (J) was determined as in (C) and ratios calculated as in (D). For statistical analysis, METs were performed. ELISA, LDH and immunoblot data are representative of at least three independent experiments. Data are depicted as mean ±SD. Significance levels are denoted as *p < 0.05, **p < 0.01, ***p < 0.001, and ns for not significant.

### NEK7 has a more pronounced role at suboptimal activating conditions

We next speculated that different intensities of the NLRP3 inflammasome activating signals might affect NEK7-dependency and carefully titrated three different NLRP3 activators (Figure 3). At lower concentrations of K^+^ efflux-dependent activators nigericin^13^, (Figure 3A, B), extracellular ATP^13^ (Figure 3C, D), or the K^+^ efflux-independent NLRP3 triggering imidazoquinoline CL097 (Figure 3E, F)^28^, not only the total amounts of IL-1β were reduced (Figure 3A, C, E), but also, the dependency on NEK7 was more pronounced (Figure 3B, D, F). The same trend was also observed for pyroptosis, again determined by LDH release assay (Figure 3A-F). Both IL-1β release and pyroptosis are facilitated by the pore-forming caspase-1 substrate GSDMD^30^. Consistent with our results for IL-1β secretion, LDH release, and caspase-1 cleavage, also the effect of NEK7 on cleavage of GSDMD was more pronounced when using lower concentrations of the three activators and became more independent at higher concentrations (Figure 3G-L). In summary, NLRP3 activation appears to be strongly NEK7-dependent under suboptimal activation conditions but becomes largely independent when optimal activator concentrations are used.

**Figure 3:**
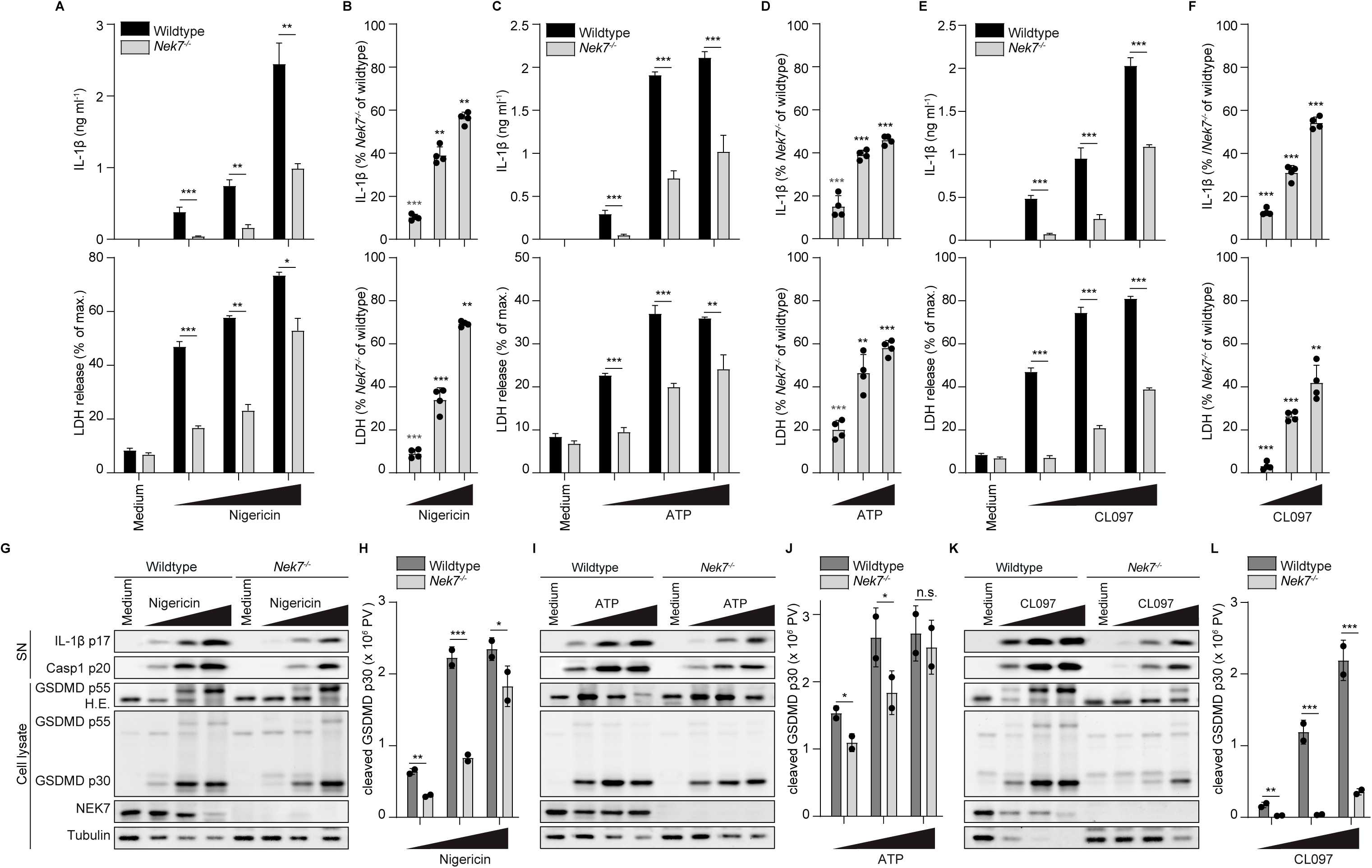
Concentration-dependent role of NEK7 in NLRP3 activation. (**A-L**) Primed BMDMs from wildtype and *Nek7^-/-^* mice stimulated with increasing concentrations of nigericin (1.25, 2.5, 5 µM; A, B, G, H), ATP (2.5, 5, 7.5 mM; C, D, I, J) or CL-097 (50, 100, 150 µM; E, F, K, L) for 2 h. (A, C, E). Release of IL-1β and LDH were determined from cell-free supernatants of technical triplicates by ELISA or a colorimetric assay, respectively. (B, D, F) Residual inflammasome activation in *Nek7^-/-^* cells is depicted as percentage relative to wildtype. Each data point is derived by calculating the ratio of the mean of three technical replicates per genotype. Data of four independent experiments, one depicted in (A, C, E). (G, I, K) Immunoblot analysis of cell lysates and cell-free supernatants from LPS-primed wildtype and *Nek7^-/-^* BMDMs. (H, J, L) Cleavage of GSDMD was quantified from immunoblots and depicted as 16-bit pixel value (PV). Samples are corrected to the respective medium control using the control protein tubulin. Data from two biological replicates, one depicted in (G, I, K). For statistical analysis, METs were performed. Data are depicted as mean ±SD. Significance levels are denoted as *p < 0.05, **p < 0.01, ***p < 0.001, and ns for not significant.

### The absence of NEK7 slows down inflammasome activation

We finally turned to the impact of the factor time on the degree for NEK7-dependency and performed time-course experiments. The influence of NEK7 on NLRP3 inflammasome activation turned out to be large at early time points but became less pronounced after longer stimulation (Figure 4A-B). We hypothesized that the observed impact of dosing (Figure 3) and of kinetics on the involvement of NEK7 might be related. This idea was triggered by our observation that – in wildtype cells – stimulation of NLRP3 with suboptimal activator doses leads to a reduced velocity of NLRP3 activation (Figure 4C). We therefore combined the two factors of time and dose, carefully titrating two different NLRP3 activators, nigericin and CL097 on wildtype and NEK7-deficient cells and monitoring inflammasome activation over time (Figure 5A, B and S2A, B). NEK7 dependency decreased both over time and with increasing activator doses. As such, IL-1β and LDH release were almost completely NEK7-dependent even for high doses of nigericin or CL097 at very early time points (0.5 h), but became completely NEK7-independent at late time points (5 h). Vice versa, even low activator doses were able to trigger substantial inflammasome activation in the absence of NEK7 at later time points. To mathematically model and evaluate the impact of NEK7 on NLRP3 activation, we performed more extensive dose-response/time course experiments, applied Hill curve fitting, and derived maximal velocity values for each activator concentration from the model (Figure 5C, D and S3C, D)^31^. Over all, NEK7-deficient cells appeared to respond up to four times slower than wildtype cells. For example, for 5 or 10 µM of nigericin, wildtype cells reach IL-1β concentrations of around 2 ng ml^-1^ already at 0.5 h, while NEK7-deficient cells need 2 h to release comparable amounts (Figure 5A).

**Figure 4:**
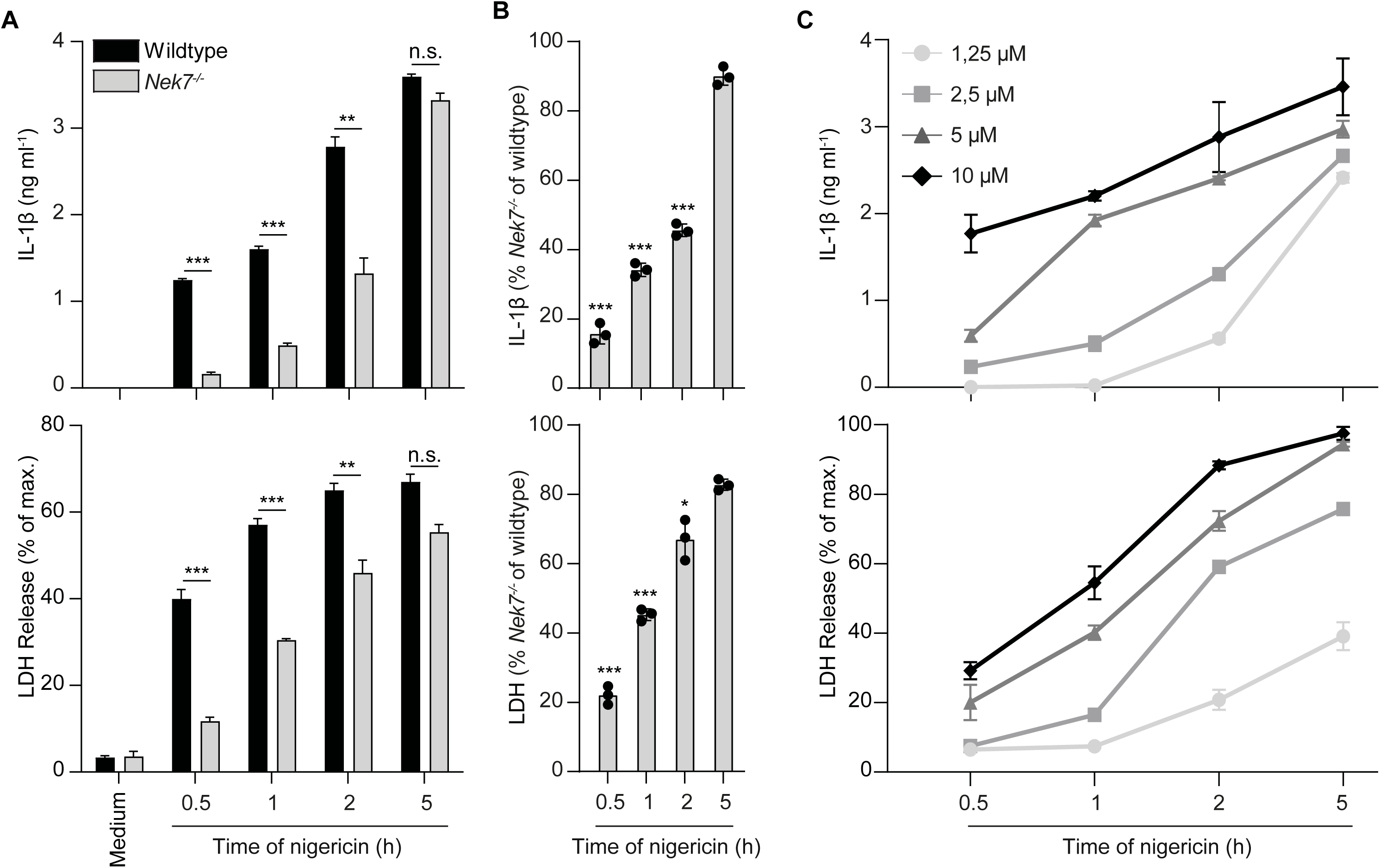
NEK7 is dispensable for NLRP3 activation at late time-points. (**A**) Primed BMDMs from wildtype and *Nek7^-/-^* mice were stimulated with 5 µM nigericin for 0.5, 1, 2, or 5 h. (**B**) Residual inflammasome activation in *Nek7^-/-^* cells is depicted as percentage relative to wildtype. Each data point is derived by calculating the ratio of the mean of three technical replicates per genotype. Data of three independent experiments, one depicted in (A). (**C**) LPS-primed BMDMs from wildtype mice were stimulated with increasing concentrations of nigericin (1.25, 2.5, 5, 10 µM) for 0.5, 1, 2 or 5 h. (A, C) Releases of IL-1β and LDH were determined from cell-free supernatants of technical triplicates by ELISA or a colorimetric assay, respectively. Result are representative of at least three independent experiments. For statistical analysis, METs were performed. Data are depicted as mean ±SD. Significance levels are denoted as *p < 0.05, **p < 0.01, ***p < 0.001, and ns for not significant.

**Figure 5:**
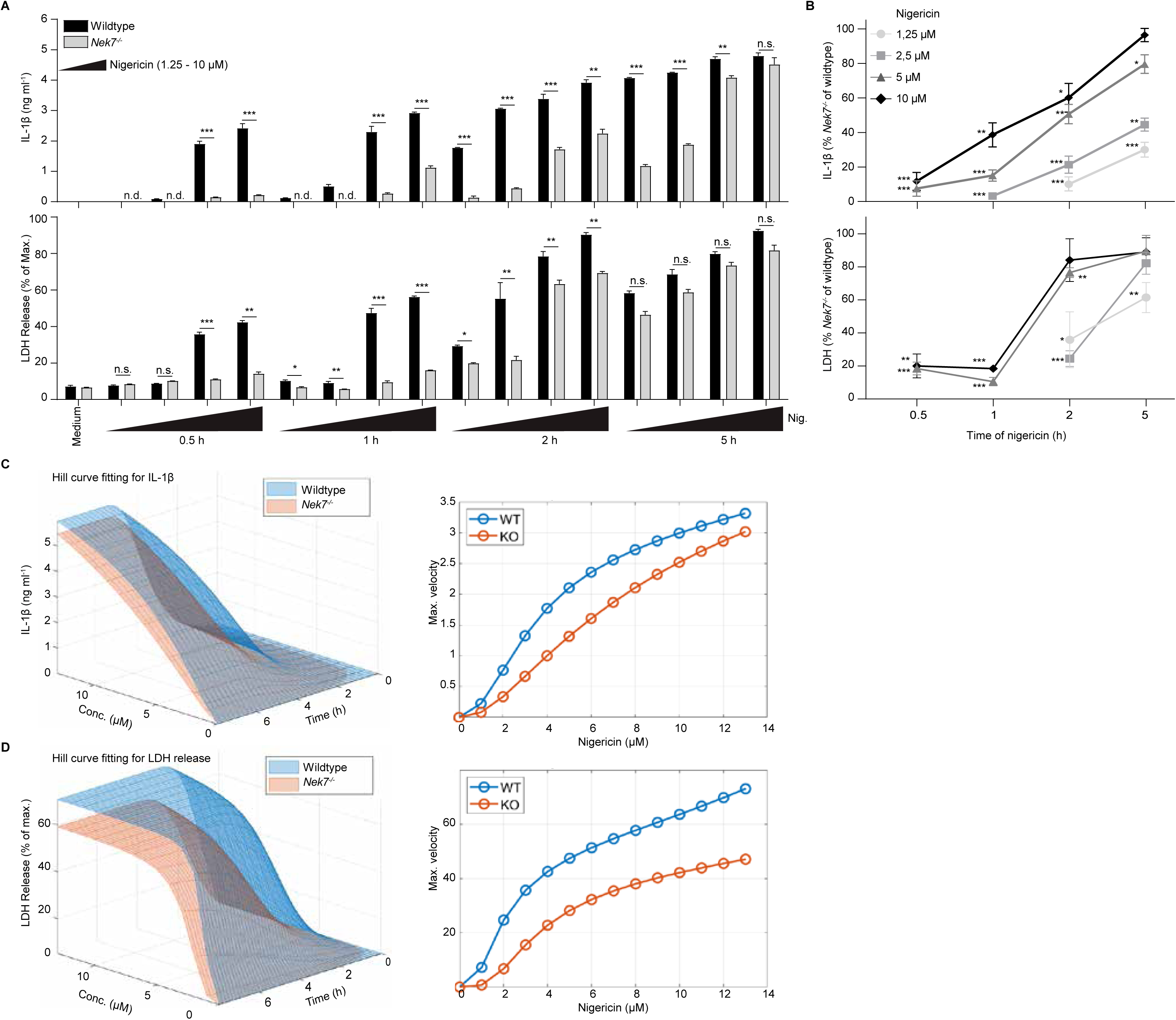
Absence of NEK7 decelerates NLRP3 inflammasome activation. (**A, B**) Primed BMDMs from wildtype and *Nek7^-/-^* mice were stimulated with 1.25, 2.5, 5, and 10 µM nigericin for 0.5, 1, 2 or 5 h. Release of IL-1β and LDH was determined from cell-free supernatants of technical triplicates by ELISA and a colorimetric assay, respectively. (B) Residual inflammasome activation in *Nek7^-/-^* cells is depicted as percentage relative to wildtype. Each data point is derived by calculating the ratio of the mean of three technical replicates per genotype. Data of three independent experiments, one depicted in (A). For statistical analysis, METs were performed. Data are depicted as mean ±SD. Significance levels are denoted as *p < 0.05, **p < 0.01, ***p < 0.001, and ns for not significant. (C, D) Time and dose data for IL-1β and LDH release, of LPS-primed wildtype and *Nek7^-/-^* BMDMs after stimulation with nigericin (1-13 µM) for (0.25-6 h) were used to generate a 3D reconstruction of individual Hill curves and derive maximal velocity values. Data are representative of three (A) and two (C, D) independent experiments.

Our data and model suggested a greater difference in velocity and also in maximal amplitude for LDH release than for IL-1β production, especially for CL097, for which the model predicted that NEK7-deficient cells would eventually overshoot wildtype cells (Figure 5C, D and S3C, D). This might reflect delayed inflammasome activation in the absence of NEK7, potentially allowing for a longer priming period for proIL-1β production, leading to higher concentrations. Indeed, under our standard priming conditions of LPS treatment for 3 h, proIL-1β concentrations have not yet reached their maximum, further increasing at later time points (Figure S3A, B). When we extended the experiments to longer time points, we could indeed observe that NEK7-deficient cells can exceed wildtype cells for IL-1β production but not for LDH release (Figure S3C, D). As the model predicted this was somewhat more pronounced for CL097 than for nigericin, likely reflecting additional priming through direct TLR7 activation by the imidazoquinoline^28^. Therefore, LDH release assay data apparently reflect the role of NEK7 more faithfully, while evaluating IL-1β values can even yield an underestimation due to further priming.

### NEK7 accelerates NLRP3 inflammasome activation on a single cell level

In order to elucidate the rate of inflammasome activation on a single cell level, we began by monitoring inflammasome-dependent pyroptosis by flow cytometry and live cell imaging (Figure 6A-D). Early membrane damage during pyroptosis allows fluorescent probes like 7AAD, Draq7, or labelled Annexin V access to the cell, where they intercalate with DNA or interact with phosphatidylinositol accessible on the inner leaflet as another consequence of membrane damage, respectively^16^. Also on a single cell level, the absence of NEK7 caused a 3-4x lower maximal rate of membrane permeabilization. In wildtype cells analyzed by flow cytometry, 50% of cells became Annexin V-positive withing 15 min, while this level was reached in the absence of NEK only after 45 min (Figure 6A, B). Similarly, when visualizing pyroptosis by microscopy, we observed a maximal rate of ∼5,8% of cells per minute to become Draq7 positive in wildtype cells, while in NEK7-deficient cells, this rate was only at ∼1%. However, in contrast to ELISA, LDH release, or flow cytometry data, in microscopy, there was no delay observed in the onset of pyroptosis with the first rare events of positive cells in both genotypes detected always at the same time point, 5-10 minutes after stimulation (Figure 6C, D). Taken together, our data to this point suggested that the presence of NEK7 increases the rate of individual cells to form an NLRP3 inflammasome by a factor of up to four.

**Figure 6:**
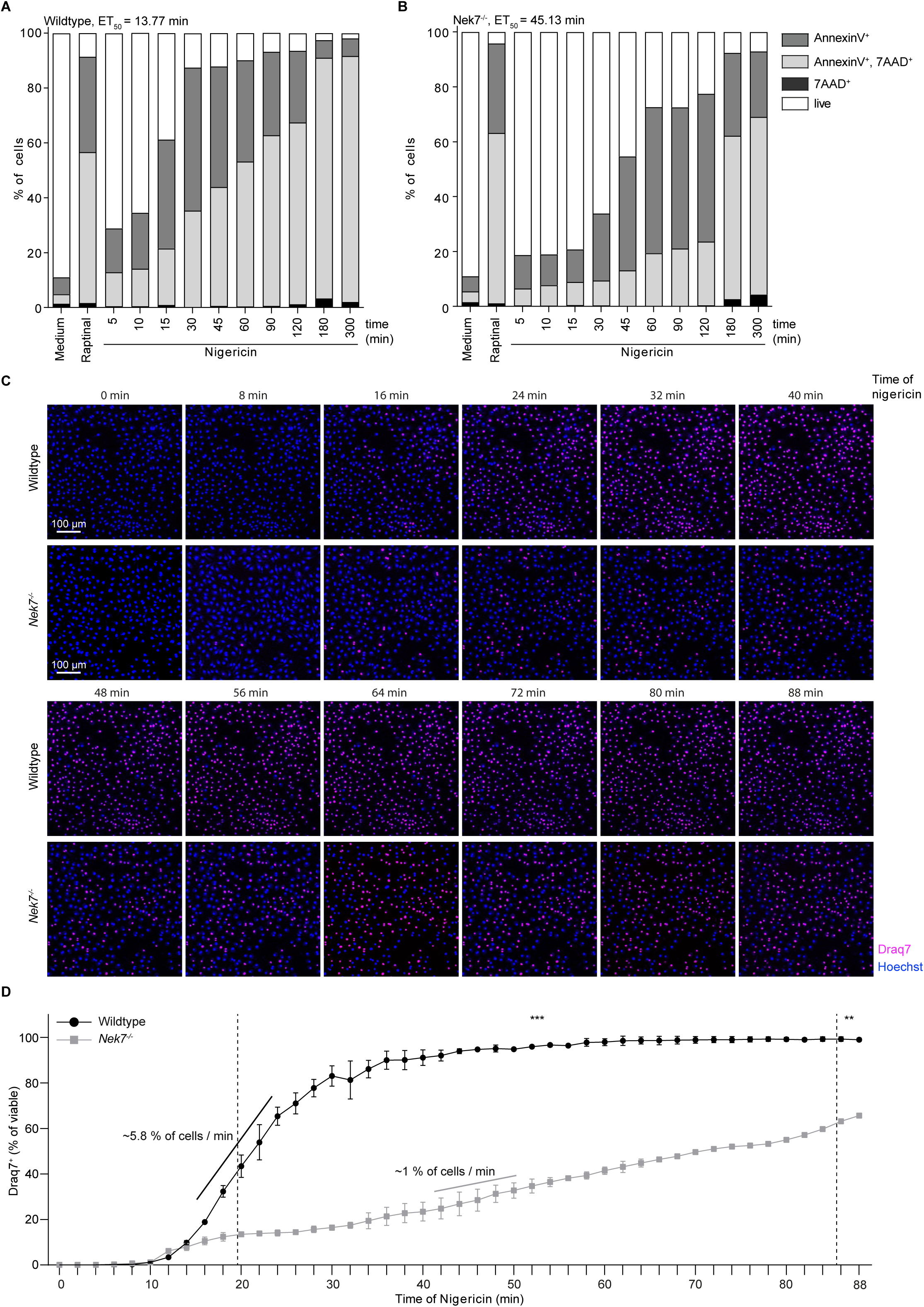
NEK7 accelerated NLRP3 inflammasome activation on a single cell level. (**A, B**) Primed BMDMs from wildtype and *Nek7^-/-^* mice were stimulated with 5 µM nigericin for intervals between 5-300 min as depicted. For comparison to apoptosis, 10 µM raptinal was applied for 90 min. Cell death was assessed by AnnexinV/7-AAD labelling and flow cytometry in wildtype cells (A) and N*ek7^-/-^* cells (B). (**C**) Primed BMDMs from wildtype and *Nek7^-/-^*mice were transferred to a live cell imaging system (controlled temperature and CO_2_ level), stained with Hoechst (blue) and Draq7 (pink), stimulated with 5 µM nigericin, and analysed by confocal fluorescence microscopy for up to 88 min. Images are representative for 3 images analyzed per condition, depicting about 390-430 cells representative for 50,000 cells per well. Scale bar represents 100 µm. Data is representative of four independent experiments (**D**) Ratio of Draq7-positive nuclei to total number of nuclei in wildtype and *Nek7^-/-^* cells was determined for each time point as depicted. Data from two independent experiments, one depicted in (C). For statistical analysis, minimal effect tests were applied. Data are depicted as mean ±SD. Significance levels are denoted as *p < 0.05, **p < 0.01, ***p < 0.001, and ns for not significant.

### Dependency of NLRP3 inhibitors on NEK7

Due to its disease-promoting activity, NLRP3 is a prominent potential therapeutic target. Sulphonylurea drugs such as MCC950/CRID3 and its derivates target NLRP3 directly and effectively, acting independent of NEK7^4,32–34^. In spite of numerous endeavors to reduce toxicity, no drugs of this family have reached the marked yet, and chemically distinct alternatives are being explored. For several reported candidates, a mode-of-action involving NEK7 has been implicated^35–38^. Our observations suggested that inhibitors that engage NEK7 or the NEK7-NLRP3 interaction should only have an incomplete inhibitory effect and should not affect the residual signal we observed in NEK7 deficiency. We revisited the contribution of NEK7 to the effect of novel NLRP3 inhibitors in murine and human cells. To avoid a potential effect of the inhibitors on priming, we added the inhibitors after 2.5 h of LPS priming and 0.5 h before addition of the inflammasome activator, and confirmed that proIL-1β and TNF production as well as cell viability were not affected under these conditions (Figure S4A-C). The endogenous metabolite itaconate and its derivative, 4-octyl itaconate (4-OI), were reported to inhibit NLRP3 by disrupting interaction with NEK7^35^. 4-OI indeed substantially reduced IL-1β and LDH release in our hands (Figure 7A-D). However, its relative inhibitory effect was not altered by NEK7-deficiency (Figure 7B, D). Another study implicated TAK1 activity in NEK7-independent inflammasome activation^38^. Although we could again confirm NLRP3 inflammasome inhibition by the TAK1 inhibitor takinib, similar to 4-OI, its relative inhibitory effect was also not reduced in NEK7-deficiency as compared to wildtype for both murine and human cells (Figure 7E-H). Similar results were also observed for the natural product alantolactone that was recently identified as an NLRP3 inhibitor and was predicted to act by interfering with NEK7 binding^37^. We could confirm inhibition of NLRP3-dependent IL-1β production by alantolactone, but found it is no less active in NEK7-deficient cells (Figure 7I-L). This suggested that 4-OI, takinib, and alantolactone can affect NLRP3 independent of NEK7. A fourth NLRP3 inhibitor reported as NEK7-dependent is entrectinib, which, as the authors found, binds directly to NEK7 and disrupts its interaction with NLRP3^36^. As – based on our data – expected for a NEK7-dependent inhibitor, entrectinib i) dampened NLRP3 only mildly in wildtype cells and ii) did not show any inhibitory effect in NEK7 knockout cells (Figure 7M-P). This confirms entrectinib as a NEK7-dependent NLRP3 inhibitor.

**Figure 7:**
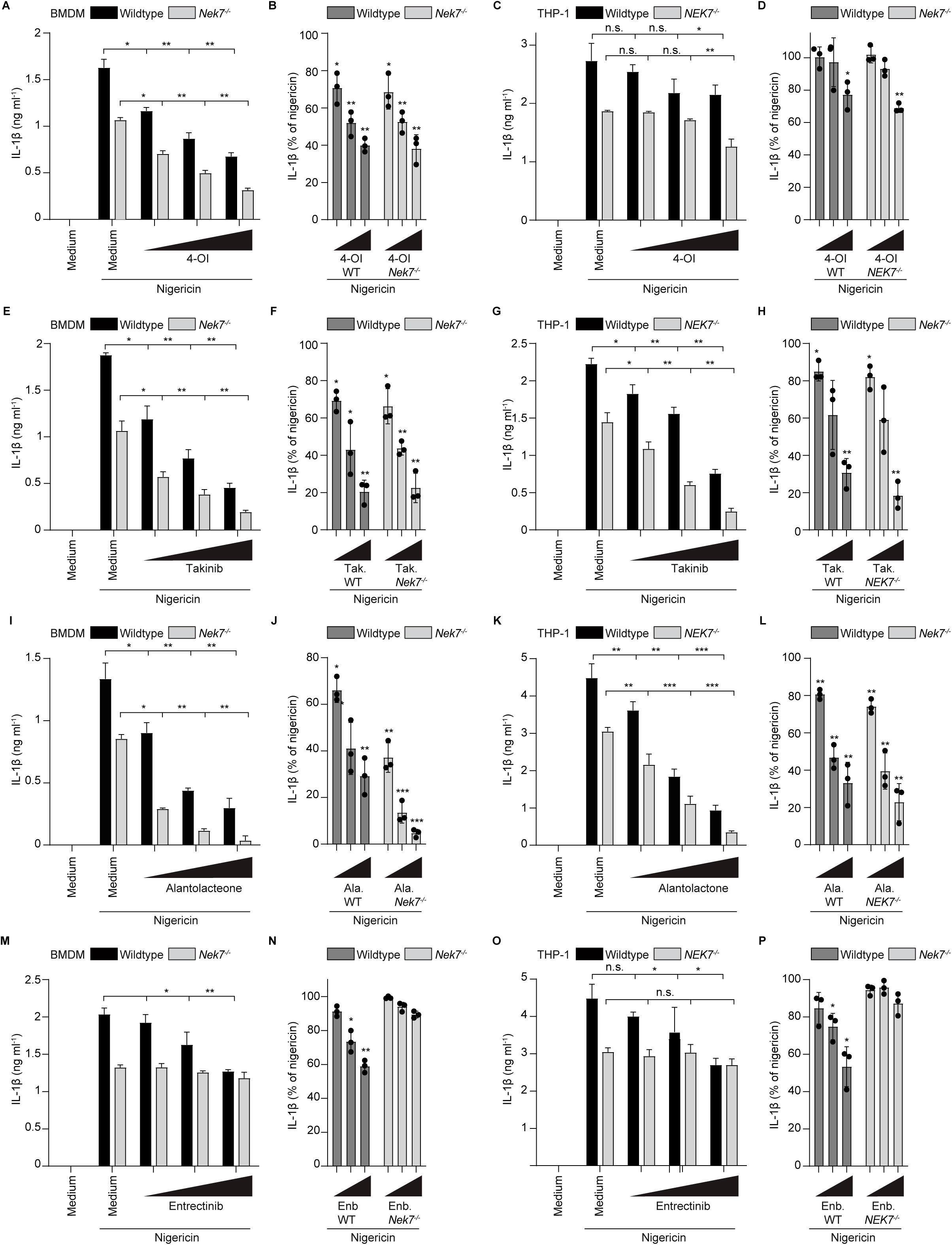
Dependency of NLRP3 inhibitors on NEK7. (**A-P**) Primed BMDMs from wildtype and *Nek7^-/-^* mice and PMA-differentiated wildtype and *NEK7^-/-^* THP-1 cells as indicated were incubated with 4-OI (250, 500, 750 µM) (A, C), Takinib (25, 50, 75 µM) (E, G), Alantolactone (2.5, 5, 10 µM) (I, K) or Entrectinib (4, 8, 16 µM) (M, O) 30 min prior to stimulation with 5 µM nigericin for 2 h. Release of IL-1β was determined from cell-free supernatants by ELISA. Data are depicted as mean ±SD of technical triplicates. The result is representative of at least three independent experiments. Data are depicted as mean ±SD. Significance levels are denoted as *p < 0.05, **p < 0.01, ***p < 0.001, and ns for not significant. (B, D, F, H, J, L, N, P) Residual inflammasome activation in *Nek7^-/-^* cells is depicted as percentage relative to wildtype. Each data point is derived by calculating the ratio of the mean of three technical replicates per genotype. Data of three independent experiments, one depicted in (A, C, E, G, I, K, M, O). For statistical analysis, one-sided T-Test was used.

## Discussion

Here we have shown that absence of NEK7 only decelerates, but cannot preclude NLRP3 inflammasome activation. The rate of inflammasome activation was up to four-fold higher in the presence of NEK7. In contrast, the eventual maximal amplitude of NLRP3 activation was not or only mildly affected. Using microscopy, we observed no delay in the onset inflammasome activation in the absence of NEK7 on the single-cell level, while batch-based read-outs (ELISA, LDH release, immunoblotting) or flow cytometry suggested a delay in the same range as the deceleration. This discrepancy is likely related to the detection threshold and background of these assays, where signals from the first few responding cells might not exceed the threshold or background, such as that wildtype cultures with a 4x increased rate of activation reach discernable levels 4x earlier that NEK7 knockout cells. Our data supports a concept in which, on the single-cell level, NLRP3 activation is a stochastic event with its likelihood being increased in the presence of NEK7.

Recent cryo-EM structures revealed that in its inactive state, NLRP3 forms *cage* or *barrel*-shaped decamers or dodecamers through interaction between different surfaces of its leucin-rich-repeat (LRR) domain^21,22^. In this conformation, the pyrin domain (PYD) engaging ASC is protected, facilitating auto-inhibition. In contrast, active NLRP3 – similar to the related inflammasome nucleator NLRC4 – forms a *wheel* decamer or undecamer^39–42^. However, in contrast to NLRC4, the LRRs of NLRP3 do not interact within the *wheel*. What proportion of the cytoplasmic pool of NLRP3 has to undergo activation to trigger inflammasome formation is unclear. Due to the prion-like activation mechanism of ASC through PYD-PYD interactions^43,44^, combined with the branching of daughter fibers involving CARD-CARD interactions^45,46^, it is imaginable that even one single active NLRP3 *wheel* complex could be sufficient to trigger the process. Although NEK7 is present in published cryo-EM structures of the active NLRP3 *wheel*, it apparently does not contribute to interactions within the *wheel*^21,22^, and our data suggests that NLRP3 can be fully activated even without NEK7. In contrast, the inactive *cage* cannot accommodate NEK7. Our data and the existing literature are consistent with a model in which NEK7 increases the chance of NLRP3 *wheel* formation by stabilizing an intermediate state, such as by serving as a kind of wedge or door-stop that prevents (re-)formation of the inactive *cage*. In this model, in the absence of NEK7, longer stimulation might either simply compensate for a decreased likelihood of a lower concentration of available active NLRP3 molecules to form the *wheel*, or more time might be needed to reach a regular cellular concentration threshold of available active NLRP3 molecules required for *wheel* formation.

Alternatively, there might be two separate NLRP3 activation mechanisms, one fast-acting and involving NEK7, and one slower and NEK7-independent. There is indeed evidence for two separate mechanisms, with an emerging study suggesting activation of *caged* NLRP3 involves vesicle binding, while uncaged NLRP3 would not require this step^27^.

What if any specific danger information NEK7 transmits towards NLRP3 remains unclear. Since NEK7 is evolutionarily more highly conserved than NLRP3^47–49^, NLRP3 might have evolved in its continued presence, and the interaction with NEK7, although accelerating NLRP3 activation, might not communicate any meaningful signaling. As such, absence of NEK7 might simply be unphysiological. However, if two separate mechanisms of NLRP3 activation with differential role of NEK7 existed, this might point towards a specific function.

NEK7 belongs to a family of cell cycle-related kinases and is enriched at the microtubule-organizing center (MTOC)^23^. Through lysine-rich basic surfaces, the inactive NLRP3 *cage* binds to acidic phospholipids generated on recycling endosomes especially when retrograde transport to the trans-Golgi network (TGN) is stalled^15^. This transport involves microtubule-dependent mechanisms, and stalled transport might cause accumulation of NLRP3 on vesicles close to the MTOC, bringing NLRP3 and NEK7 into proximity. Indeed, specifically the *caged* form of NLRP3 was shown to travel on vesicles towards the MTOC^22^. Binding of NLRP3 to phospholipids, specifically PIP, on vesicles was suggested to support activation, but our recently-published data suggested it is not sufficient^14,16^. NLRP3 binds to various phospholipids with potentially insufficient specificity to serve as a danger signal. Binding to phospholipids on vesicles might cooperate with the presence of NEK7 to increase signal specificity and open the inactive NLRP3 *cage*, while binding to other phospholipids on other membranes in the absence of elevated local NEK7 concentrations might be insufficient to sufficiently open up the *cage*. In such a scenario, NEK7 would only be required for activation of caged NLRP3, while the hypothesized cage-independent activation mechanism would also not involve NEK7^27^. Further research is required to elucidate whether indeed, two pathways of NLRP3 activation with differential requirements for NEK7 exist. Entrectinib, specifically blocking NEK7-dependent NLRP3 activation by directly binding to NEK7 at the site of NLRP3 interaction, will be useful for such research^36^.

Our findings suggest that therapeutic NLRP3 inhibitors that target the interaction with NEK7 would only have a partial effect. This might be favorable to limit the immunosuppressive effect. Inhibiting NLRP3 itself is generally desirable to leave the protective effects of the other inflammasomes and of IL-1 intact^50^. There is no consensus yet concerning the beneficial physiological function of NLRP3 that justifies its evolutionary conservation in spite of its disease-promoting activities. However, it is triggered by specific pathogens including viruses and fungi, and prolonged NLRP3 inhibition might increase the risk of infection with these^16,51^. Targeting NEK7 for NLRP3 inhibition might allow to hit a sweet spot of sufficiently dampening its activity to achieve a therapeutic effect without severely impeding host defense.

Although some researchers reported that NEK7 deficiency is lethal in mice, their initial discovery through a murine mutagenesis screen and the subsequent generation of knockout mice by CRISPR-Cas9-based targeted gene disruption reported by the same authors suggests that NEK7 deficiency does not necessarily affect viability^38,52^. Therefore, cell and animal tools are available to test an involvement of NEK7 in the effect of newly discovered and already reported NLRP3 inhibitors and thereby elucidate their mode-of-action and estimate their maximal potency. This will promote the development of approaches to tune the degree of NLRP3 inhibition for precision medicine.

## Acknowledgements

We thank Romeo Ricci for providing NLRP3- and NEK7-deficient THP-1 cells, and the Lighthouse Core Facility, funded in part by the Medical Faculty, University of Freiburg (Project Numbers 2023/A2-Fol; 2021/B3-Fol) and the DFG (Project Number 450392965) for the providence of microscopy imaging systems including the Zeiss CellDiscoverer7 w/LSM900 (Project Number 452993504) and technical support by Franck A. Ditengou and Marie Follo.

## Author Contributions

S.W., T.C., C.D., E.N., F.K., A.K., O. Gorka, and C.J.G. performed experiments and analyzed data. C.K., performed mathematical modelling. S.W. prepared figures and wrote figure legends and the methods section of the manuscript with input from other authors. O. Groß and C.J.G. wrote the main text of the manuscript. O. Groß conceived and oversaw the project.

## Figure Legends

**Figure S1.**
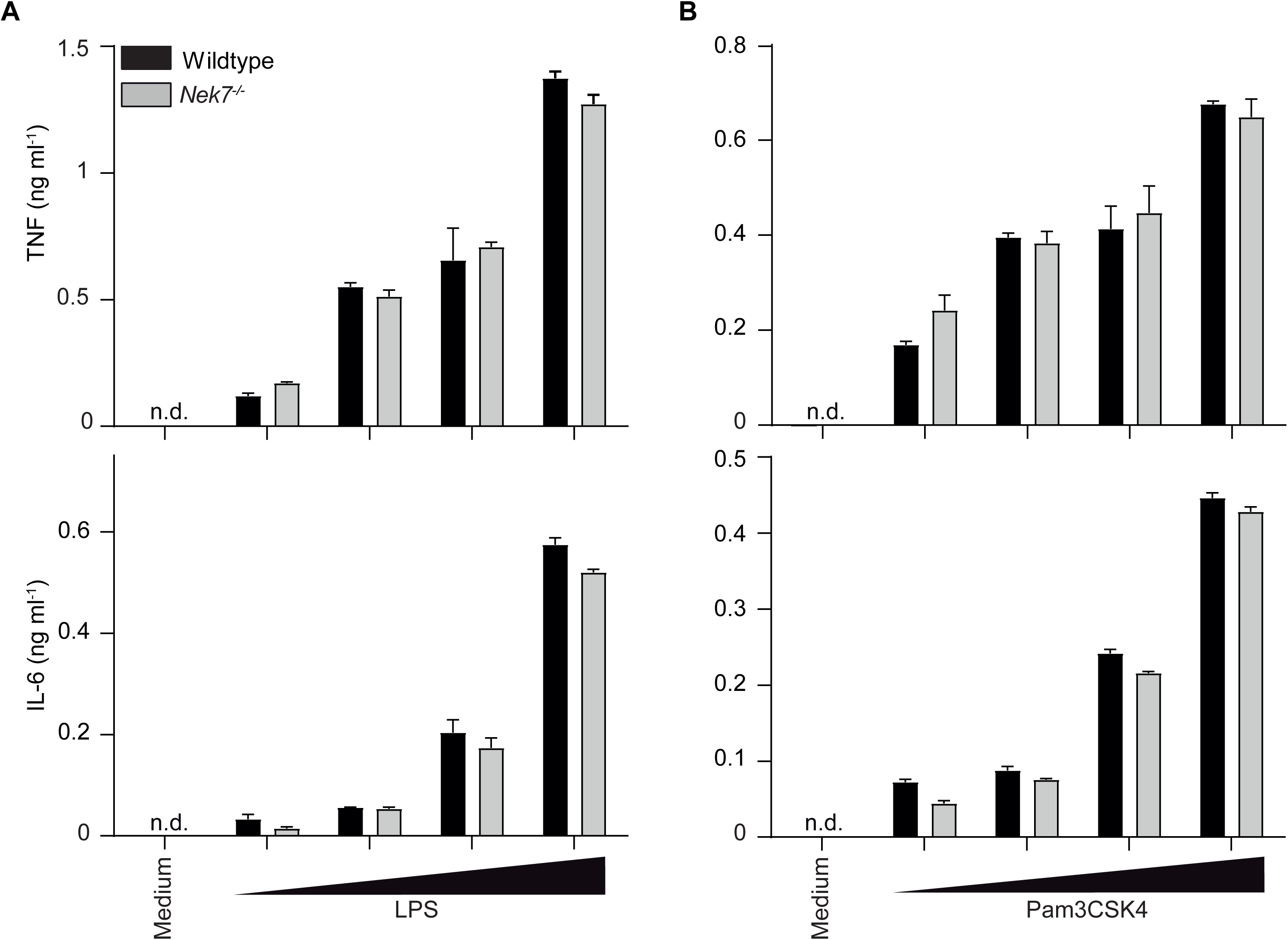
corresponding to main Figure 2: Priming does not affect the influence of NEK7 on NLRP3. (**A, B**) BMDMs from wildtype and *Nek7^-/-^* mice were primed with increasing concentration of LPS (0.4, 2, 10, 50 ng ml^-1^; A) or Pam3CSK4 (0.125, 0.25, 0.5, 1 µg ml^-1^; B) for 3 h. TNF and IL-6 release were determined from cell-free supernatants by ELISA, N = 3 technical replicates. The result from each experiment in this figure is representative of at least three independent experiments. Data are depicted as mean ±SD.

**Figure S2.**
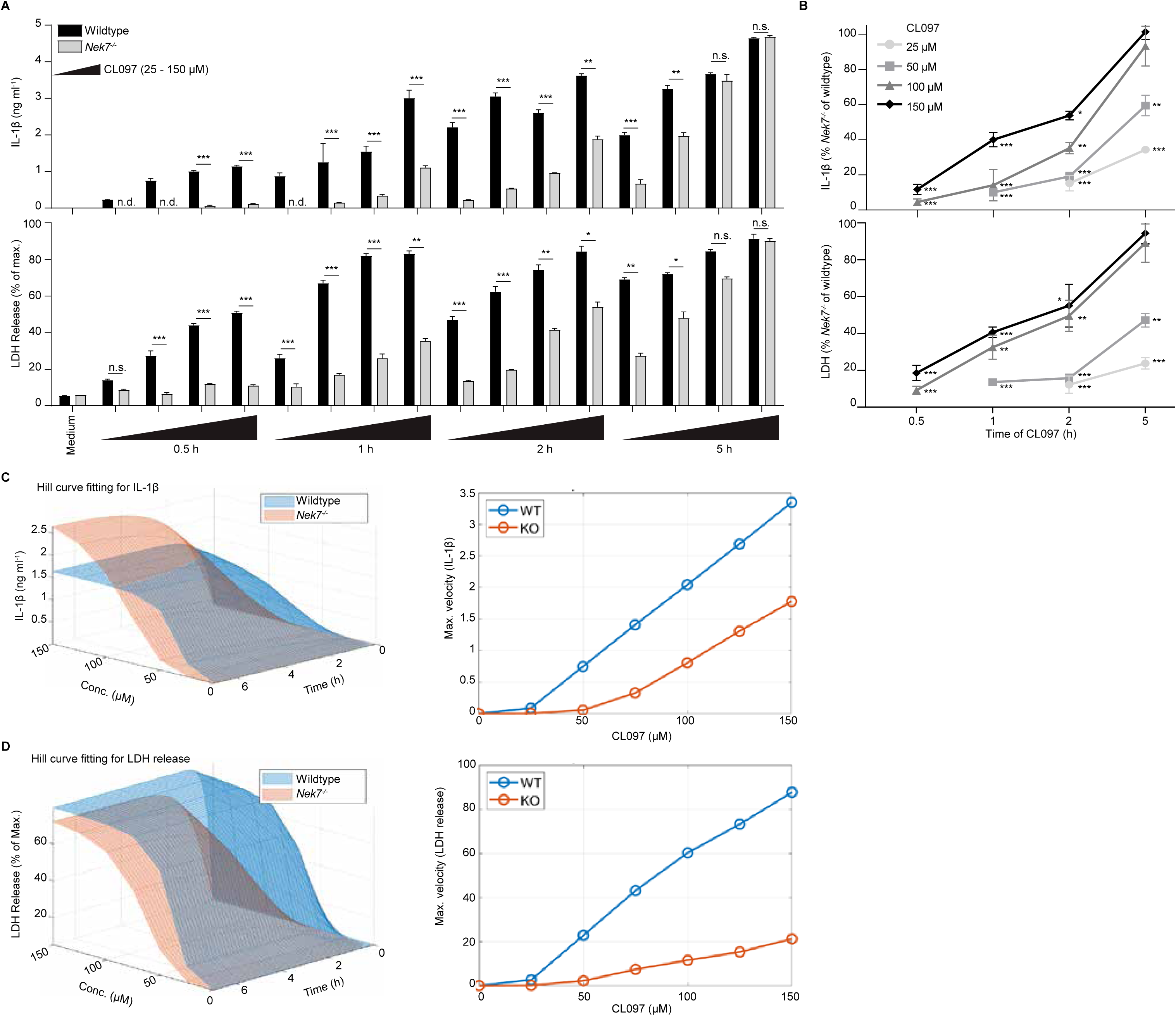
corresponding to main Figure 5: Absence of NEK7 decelerates NLRP3 inflammasome activation. (**A, B**) Primed BMDMs from wildtype and *Nek7^-/-^*mice were stimulated with 25, 50, 100 and 150 µM CL097 for 0.5, 1, 2 or 5 h. Release of IL-1β and LDH was determined from cell-free supernatants of technical triplicates by ELISA and a colorimetric assay, respectively. (B) Residual inflammasome activation in *Nek7^-/-^* cells is depicted as percentage relative to wildtype. Each data point is derived by calculating the ratio of the mean of three technical replicates per genotype. Data of three independent experiments, one depicted in (A). For statistical analysis, METs were performed. Data are depicted as mean ±SD. Significance levels are denoted as *p < 0.05, **p < 0.01, ***p < 0.001, and ns for not significant. (**C, D**) Time and dose data for IL-1β and LDH release, of LPS-primed wildtype and *Nek7^-/-^* BMDMs after stimulation with CL097 (25-150 µM) for (0.5-6 h) were used to generate a 3D reconstruction of individual Hill curves and derive maximal velocity values. Data are representative of three (A) and two (C, D) independent experiments.

**Figure S3.**
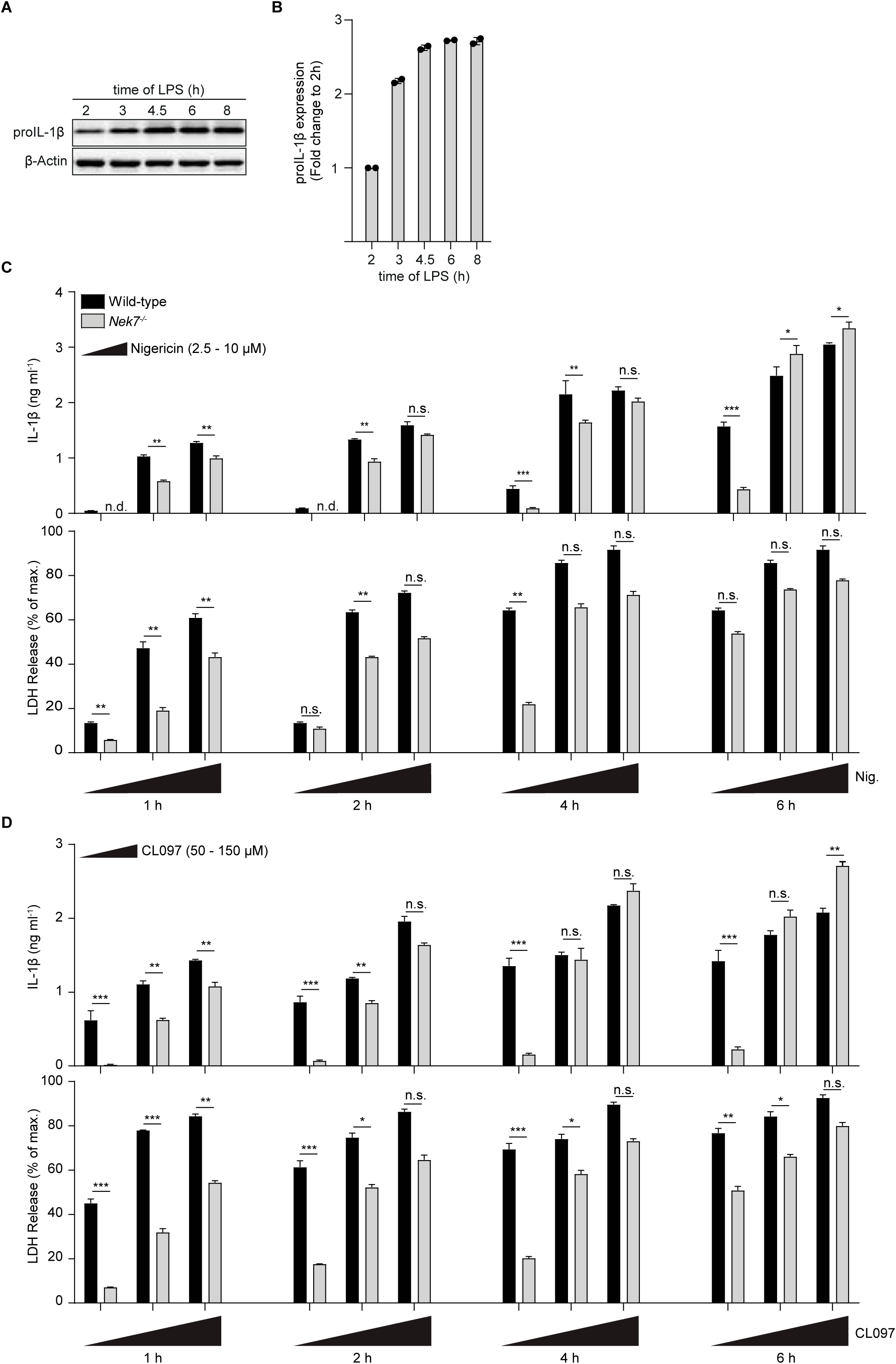
corresponding to main Figure 5: IL-1β production from NEK7-deficient cells can exceed that from wildtype cells at late time points. (**A, B**) BMDMs from wildtype mice were primed with 20 ng ml^-1^ LPS for 2, 3, 4.5, 6 and 8 h, production of proIL-1β was determined by immunoblotting (A), and quantified from immunoblots and depicted as fold change compared to the shortest time-point. Samples are corrected to the respective control signal for β-actin (B). (**C, D**) BMDMs from wildtype and *Nek7^-/-^* mice were primed with 20 ng ml^-1^ LPS and stimulated with increasing concentrations of nigericin (2.5, 5, 10 µM; C) or CL097 (50, 100, 150 µM; D) for 1, 2, 4, or 6 h. Releases of IL-1β and LDH were determined from cell-free supernatants of technical triplicates by ELISA and a colorimetric assay, respectively. The result from A and B are representative of at two and C and D of at least three independent experiments. For statistical analysis, minimal effect tests were applied. Data are depicted as mean ±SD. Significance levels are denoted as *p < 0.05, **p < 0.01, ***p < 0.001, and ns for not significant.

**Figure S4.**
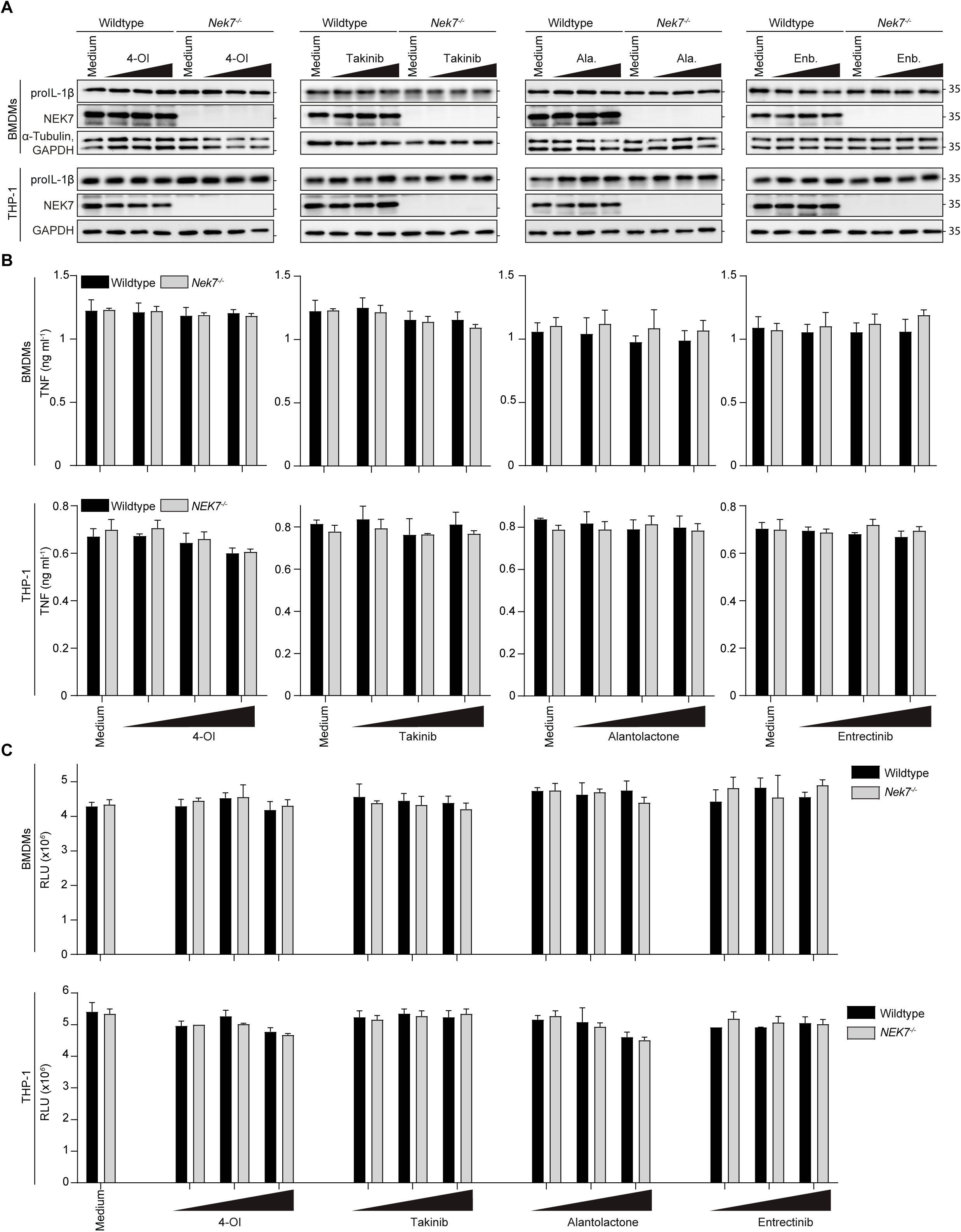
corresponding to main Figure 7: Dependency of NLRP3 inhibitors on NEK7. (**A-C**) Primed BMDMs from wildtype and *Nek7^-/-^* mice and PMA-differentiated Wildtype and *NEK7^-/-^* THP-1 were incubated with 4-OI (250, 500, 750 µM), Takinib (25, 50, 75 µM), Alantolactone (2.5, 5, 10 µM) or Entrectinib (4, 8, 16 µM) for 30 min. Production of proIL-1β was determined by immunoblotting (A) and release of TNF was determined from cell-free supernatants of technical triplicates by ELISA (B). Cell viability was assed 2 h after addition of the inhibitors by determining cellular ATP levels form technical triplicates using a luminescent assay (C). Data are representative of at least two independent experiments and are depicted as mean ±SD.

## Materials and Methods

### Mice

*Nlrp3^-/-^*^29^, *Pycard^-/-^*^53^, *Casp1^-/-^*^54^ (25) and *Nek7^-/-^*^55^ and wildtype mice on C57BL/6 background were housed under SOPF or SPF conditions at the Center for Experimental Models and Transgenic Services (CEMT, Freiburg, Germany), the Zentrum für Präklinische Forschung (ZPF, Munich, Germany), or Charles River Laboratories (Calco, Italy) in accordance with local guidelines.

### Cell lines

Cell lines were cultured in T75 flasks at 37 °C, 5% CO_2_ in a humidified incubator and continuously passaged according to the cell density to prevent overgrowth. Media were typically supplemented with 10 % fetal calf serum (FCS) (Gibco) and 100 U ml^-1^ penicillin-streptomycin (Gibco). THP-1 cells (RRID:CVCL 0006) were cultured in RPMI (Gibco). For inflammasome stimulation, the cells were differentiated for 3h with 100 ng ml^-1^ phorbol 12-myristate 13-acetate (PMA) and rested for at least 12 h^56^.

### BMDCs and BMDMs Preparation and Stimulation

Cells were cultured at 37 °C, 5% CO2 in a humidified incubator. Murine bone marrow-derived macrophages (BMDMs) were differentiated from tibial and femoral bone marrow as previously described in detail^57,58^. Recombinant human M-CSF was from Immunotools and was used at 100 ng ml^-1^. After 6 - 8 days of differentiation, cells were plated in 96-well plates at a density of 0.8 - 1x10^5^ cells per well, primed with 5 - 50 ng ml^-1^ E. coli K12 ultra-pure LPS (InvivoGen) for 1 - 3 h (typically 50 ng ml-1 for 3h) or 0.1 - 1 µg ml^-1^ Pam3CSK4 (InvivoGen) for 1-3 h (typically 1 µg ml^-1^ for 3 h). Afterward priming, cells were treated with inflammasome activators for 0.5 - 6 h. Inflammasome activators and stimuli were used as follows: 2.5, 5, 7.5 mM ATP (Sigma), 1.25, 2.5, 5 µM nigericin (Sigma), 300 μg ml-1 MSU (prepared as previously described^57^), 1 μg ml-1 poly(dA:dT) (InvivoGen) (transfected with Lipofectamine 2000, Invitrogen). All stimulations were performed in triplicates and cytokine production in cell-free supernatants was measured by ELISA, as previously described ^57,58,59^. Inhibitors for NLRP3 inflammasome activation were added after 2.5 h priming and 30 min before the activator. Alantactone (Hycultec GmbH) was used at 2.5-10 µM, Entrectinib (Targetmol) at 4-16 µM, 4-octyl itaconate (Sigma-Aldrich) at 250 to 750 µM, and Takinib (Sigma-Aldrich) at 25-75 µM.

### Immunodetection of Proteins

For cytokine quantification of cell-free supernatants, ELISA kits for murine IL-1β, IL-6 and TNF (Thermo, R&D Systems) were used. ELISA data is depicted as mean ±SD of technical triplicates as previously described^57,58^. For immunoblotting cell-free supernatants were mixed with 3× Tschopp-buffer (187.5 mM Tris–HCl, pH 6.8, 6% w/v SDS, 0.03% w/v Phenol Red, 30% w/v Glycerol) and cells lysed in 1× Tschopp-buffer. Triplicates were pooled and proteins were separated by SDS-PAGE and transferred to nitrocellulose membranes using standard techniques^57,58^. The following primary antibodies were used: anti-Caspase-1 (p20) mAb (Casper-1) (Adipogen), IL-1β/IL-1F2 pAb (R&D Systems), anti-GSDMD (EPR19828) (Abcam), anti-NLRP3/NALP3 mAb (Cryo-2) (Adipogen), anti-Nek7 mAb (EPR4900) (Abcam) and tubulin mAb (T5168) (Sigma-Aldrich). Signal intensities were determined by calculating the mean 16-bit grey pixel value (PV) of individual bands using Photoshop (Adobe Version 11). For each lanes equal areas were determined including all of the signal and a minimal amount of background. An equal empty area was measured to subtract the background from all signal measurements.

### Cell Viability Assays

Lytic cell death was determined by measuring LDH release from cell-free supernatants using a colorimetric assay (Promega, Takara) according to the manufacturer’s protocol. Medium served as blank value and was subtracted from the sample values. Results were plotted as percentage of 100 % dead cells which was determined by lysing cells with lysis buffer 45 min prior to collection of the cell supernatants. Intracellular ATP was quantified using the CellTiter-Glo Luminescent Cell Viability Assay (Promega) according to the manufacturer’s protocol. Luminescence was recorded and depicted as relative light units (RLU). Data is depicted as mean ±SD of technical triplicates.

### Fluorescence Imaging

BMDMs were plated at a density of 0.5 – 1x10^5^ cells per well in 8-well µ-slides (IbiTreat, Ibidi). Cells were primed with 50 ng ml^-1^ LPS for 2 h followed by treatment with 5 µM nigericin. Cells were washed with PBS, fixed in 4 % paraformaldehyde (PFA) for 10 min and permeabilized in PBS with 0.1 % (v/v) Triton X-100 for 5 min. Cells were stained with anti-ASC primary antibody (AL177, Adipogen) diluted in blocking buffer consisting of PBS, 5 % FCS and 0.1 % Triton X-100, followed by anti-rabbit IgG cross-absorbed secondary antibody (Alexa Fluor 555) (Invitrogen), and finally mounted in Vectashield antifade mounting medium containing DAPI (Vector Laboratories).

For live cell imaging at 37 °C in a 5 % CO2, humidified atmosphere, the cells were primed with 50 ng ml^-1^ LPS for 2 h and then stained as indicated. Draq7 (BioLegend) and Hoechst 3342 (Invitrogen) were used to stain dead cells and nuclei, respectively.

Confocal microscopy was performed with a Leica SP8 confocal microscope equipped with a 63×/1.40 and 40×/ 1.25 oil objective (Leica Microsystems) keeping the laser settings of the images constant to allow direct comparison of signal intensities between images of the same channel within the same experiment.

### Cell death characterization by flow cytometry

Pacific Blue Annexin V Apoptosis Detection Kit with 7-AAD (BioLegend) was used to characterize cell death by flow cytometry. To this end, BMDMs were treated with nigericin as indicated, harvested with HBSS/EDTA, and transferred to 96-well V-bottom plates. Cells were stained with Pacific Blue Annexin V and 7-AAD in Annexin V binding buffer according to the manufacturer’s protocol. The cells were washed and analyzed with a BD FACS Canto II (BD Biosciences) flow cytometer. Data were acquired with DIVA (BD Biosciences) and were analyzed with FlowJo software.

### Statistical analysis

Sample size was based on previous studies, not predetermined by a statistical method. For continuous observations, an unpaired t-test, or a minimal effect test (MET) was applied to test for significant effects, as indicated in the figure legends. Two-tailed t-tests were applied to test the null hypothesis. METs were performed with a 20% difference of the larger mean as threshold and responses that did significantly reject the hypothesis of such a minimal difference were considered as essentially intact. Statistical significance is indicated with asterisks: *p < 0.05; **p < 0.01; ***p < 0.001; and n.s., not significant.

### Mathematical Modelling

For mathematical modelling, the time and dose dependencies of LDH and IL-1β release, the so-called dose-dependent retarded transient function approach has been applied^31^. Within this approach, the kinetics are described by a combination of exponential functions and a transformation of the time axis to describe delayed response. Moreover, the dose dependency of the kinetic is described by Hill functions. Since we only observe a monotonic increase of both readouts over time, the kinetics only required the sustained part 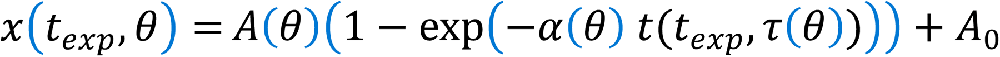. Here θ denotes all fitted parameters, *A*_0_ is the measured baseline level, *t*_*exp*_ denotes the time point of the measurement, and *t*(*t_exp_*, τ) is a nonlinear transformation of the time axis with a delay parameter τ. For mathematical details, we refer to Rachel et al.^31^. In brief, the delay acts like a time shift on the log-log scale which – in contrast to a classical delay - enables a smooth onset of the response. The dose-dependencies of *A*, α, τ are introduced by three individual Hill functions 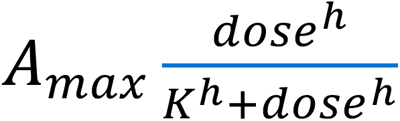, where *A_max_* denotesthe maximal response, the exponent *h* is the Hill-coefficient controlling sigmoidality and *K* is the half-maximal dose. Moreover, to account for differences between wildtype and knockout, fold-change parameters were introduced and fitted for the maximal response *A_max_* and the half-maximal doses *K*. Altogether, 15 parameters θ were fitted (1 baseline level *A*_0_, 3 half maximal doses *K* for *A*, α, τ in WT, 3 maximal responses *A_max_* for *A*, α, τ in WT, 3 Hill-coefficients *h* for *A*, α, τ in WT, as well as 4 fold factors for *K* and *A_max_* for *A*, α for the KO, and 1 error parameter σ for the residual standard deviation). These 15 parameters were jointly estimated using maximum likelihood on all data points measured for all doses and time points for WT and KO. Maximum likelihood parameter estimation was performed using multi-start optimization, jointly estimating location and error parameters^61^. Uncertainty analyses were conducted using the profile likelihood approach^61^. All these modelling analyses were performed using the Data2Dynamics modelling framework^62^ that runs under Matlab (R2021) and is available on https://github.com/Data2Dynamics. More technical and mathematical background and details are provided in Rachel *et al.*^31^ for a similar data example. For the left panels in Figure 5 C, D and Fig S2 C, D the fitted time and dose dependencies were plotted. For calculating the maximal velocity shown on the right, the maximal slope of the kinetic for each depicted dose is calculated based on difference quotients calculated from 1,000 equidistant points along the dose axes.

